# Longitudinal, Non-Invasive Imaging of the Developing Chick Heart, Vasculature, and Chorioallantoic Membrane (CAM)

**DOI:** 10.64898/2026.09.22.753481

**Authors:** Zijia Jin, Sudhanshu Kakkar, Dhaivik Chenemilla, Carol Readhead, Marianne Bronner, Simon Mahler

## Abstract

The chick embryo is a widely used vertebrate model for developmental biology, cardiovascular, and bioengineering research. We present an automated laser speckle contrast imaging (LSCI) platform for non-invasive, longitudinal, label-free imaging of the developing heart and extraembryonic vasculature of the chick embryo through the intact eggshell. The platform can detect blood vessels as small as 70 ± 17 μm in diameter and enable repeated imaging from day 0 to day 8 of incubation without disturbing normal embryonic development. It provides both structural (a map of growing extraembryonic blood vessels) and functional (blood flow dynamics and heart rate) information. Using this system, we generated the first continuous, non-invasive, time-lapse recordings of the early functioning chick heart and its extraembryonic vessels. Longitudinal imaging was performed on 15 embryos from different chicken breeds with eggshell colors white, brown and green. Analysis revealed slight variability in early cardiac and vascular morphogenesis across breeds and shell types and shows its potential for automated developmental staging. Finally, for the first time, the growing vasculature of the yolk sac and the chorioallantoic (CAM) membrane was imaged from day 4 to 8. The heart rate and growth of the extraembryonic blood vessels were quantified automatically during imaging.

## 1. Introduction

Avian models play a crucial role in advancing our understanding of disease pathophysiology and provide an essential platform for evaluating novel medical devices and therapeutics, particularly due to having a cardiovascular system similar to humans ^[1–3]^. For example, the development of the four chambered heart is similar in all amniotes^[3]^, but imaging the developing human heart under normal physiological conditions remains technically challenging. The chick embryo has been a valuable model for cardiovascular studies because of its accessibility and its rich network of extraembryonic blood vessels^[4–6]^, and close-recapitulate key aspects of human cardiac development^[1,2]^. The chick heart undergoes a well-defined developmental process characterized by specific stages. At approximately 44 hours of incubation, the heart, which is a simple tube, begins to beat^[7,8]^. It then undergoes key intermediate stages, including looping of the heart tube at around 55 hours of incubation^[9]^ and the formation of atrial and ventricular chambers by 72 hours of incubation. Cardiac neural crest cells migrate to the heart and contribute to a number of structures including septation of the outflow tract^[10]^. The heart reaches its final form as a four-chambered structure around 120 hours of incubation^[2,11]^.

The chicken embryo has an extensive extra-embryonic vasculature in the yolk sac and chorioallantoic membrane (CAM). These highly vascularized membranes perform multiple functions during embryonic development such as gas exchange and nutrient transport. The CAM has a dense capillary network which has been used to study normal and tumor angiogenesis *in vitro and vivo*^[12]^. However, noninvasive longitudinal imaging of this vasculature without perturbing normal development remains a significant challenge.

While there exist several imaging techniques including ultrasound imaging^[13]^, X-rays computed tomography^[14,15]^, and fluorescence or confocal microscopy^[16–18]^, these require windowing the shell, which partially alters natural development and shortens the live imaging time. Magnetic resonance imaging (MRI) offers non-invasive imaging^[16,18]^, but is impractical due to its high cost, complexity, and limited availability of instruments. Other non-invasive imaging modalities include fluorescence, confocal microscopy, and optical coherence tomography; however, these systems cannot penetrate the eggshell effectively and therefore the shell has to be windowed for imaging or the embryo must be grown *in vitro* ^[16–18]^.

Egg candling (also known as brightfield imaging) enables optical visualization of the chick embryo without breaking the shell, by exploiting the differences in light absorption^[19,20]^. However, its performance is susceptible to undesired external sources of light absorption, such as eggshell cracks, eggshell pigmentation, or air bubbles between the inner and outer shell membranes^[21]^. Consequently, egg candling is ineffective for imaging brown chicken eggs or quail eggs^[19,21]^. Furthermore, it only provides structural information and lacks functional imaging capabilities (blood flow and heart rate monitoring). Because inactive blood vessels and deceased embryos continue to absorb light, nonviable embryos can appear indistinguishable from living ones. These limitations confound developmental studies, making egg candling unreliable for accurately assessing embryonic viability, heart function and vasculogenesis.

Laser speckle contrast imaging (LSCI) has recently emerged as an effective technique for non-invasive imaging of the developing chick embryo’s vasculature and blood flow^[21–24]^. LSCI imaging is based on the scattering of coherent laser light by biological tissue, where the motion of red blood cells produces temporal fluctuations in the speckle pattern that enable visualization of blood flow. Blood vessels as small as 100 μm in diameter have been visualized with a recent LSCI system^[21]^. The temporal resolution of LSCI is determined primarily by the frame rate of the detector (camera) and can exceed 80 Hz. Unlike egg candling, LSCI is compatible with eggs of any shell color, including brown and quail eggs, and remains effective even in the presence of shell pigmentation, staining, or minor cracks^[21]^. In addition, LSCI provides real-time functional imaging of embryonic blood flow, enabling testing factors using image-guided injections^[23]^. Finally, LSCI is well suited for large sample studies as its automated imaging workflow enables rapid data acquisition, as showed in^[24]^ where more than 1,250 eggs were imaged in less than half a year.

In this study, we present an automated laser speckle contrast imaging (LSCI) system specifically designed for longitudinal, non-invasive and quantitative imaging of the developing chick (and other avian) heart and extraembryonic vasculature. A key advantage of the system is its ability to image embryos repeatedly throughout development without disturbing their natural incubation environment, enabling visualization and quantification of cardiac and extraembryonic vascular development at any user-defined time points from day 0 to 8 of incubation (∼38% of the 21-day incubation period).

First, we validated the performance of the automated LSCI system by comparing its measurements of both major blood vessels and capillaries with microscopy images acquired after opening the eggshell. Second, we performed longitudinal imaging (every 30 minutes) on 15 chicken eggs representing three shell types: brown (Rhode Island), white (White Leghorn), and green (Easter Egger) eggs. To the best of our knowledge, this is the first study to report a continuous, non-invasive time-lapse functional imaging of chick heart, extraembryonic yolks sac and CAM blood vessel as they develop. Cardiac heart rate and vascular morphogenesis growth rate were quantified on N=15 embryos and observed slight-to-no variation between embryos from different breeds of chicks. Finally, we report, for the first time, the non-invasive imaging of the chorioallantoic membrane (CAM) vasculature onset and development. We believe that this automated LSCI platform will open new avenues for research in heart development, developmental biology, and vasculogenesis. It can be used for quantitative testing of pharmaceutical or environmental agents that affect heart formation and function and provide new insights for congenital heart disease.

## 2. Conclusion

### 2.1 Validation of the LSCI System for Blood Vessel Imaging

The automated LSCI system (**Figure** 1A) for imaging the embryonic heart and its surrounding vasculature is shown in Figure 1A. See Methods for more details on LSCI imaging, and the experimental arrangement of our system. In Figure 1B, part of the eggshell was removed to expose the embryo for verification. The embryo was then imaged using a stereo telescope under brightfield illumination, for a high-resolution reference of the embryo and its external vasculature. A typical LSCI image of the extraembryonic blood vessels is shown in Figure 1C together with opened egg image in Figure 1D, and superimposition (Figure 1E).

**Figure 1.**
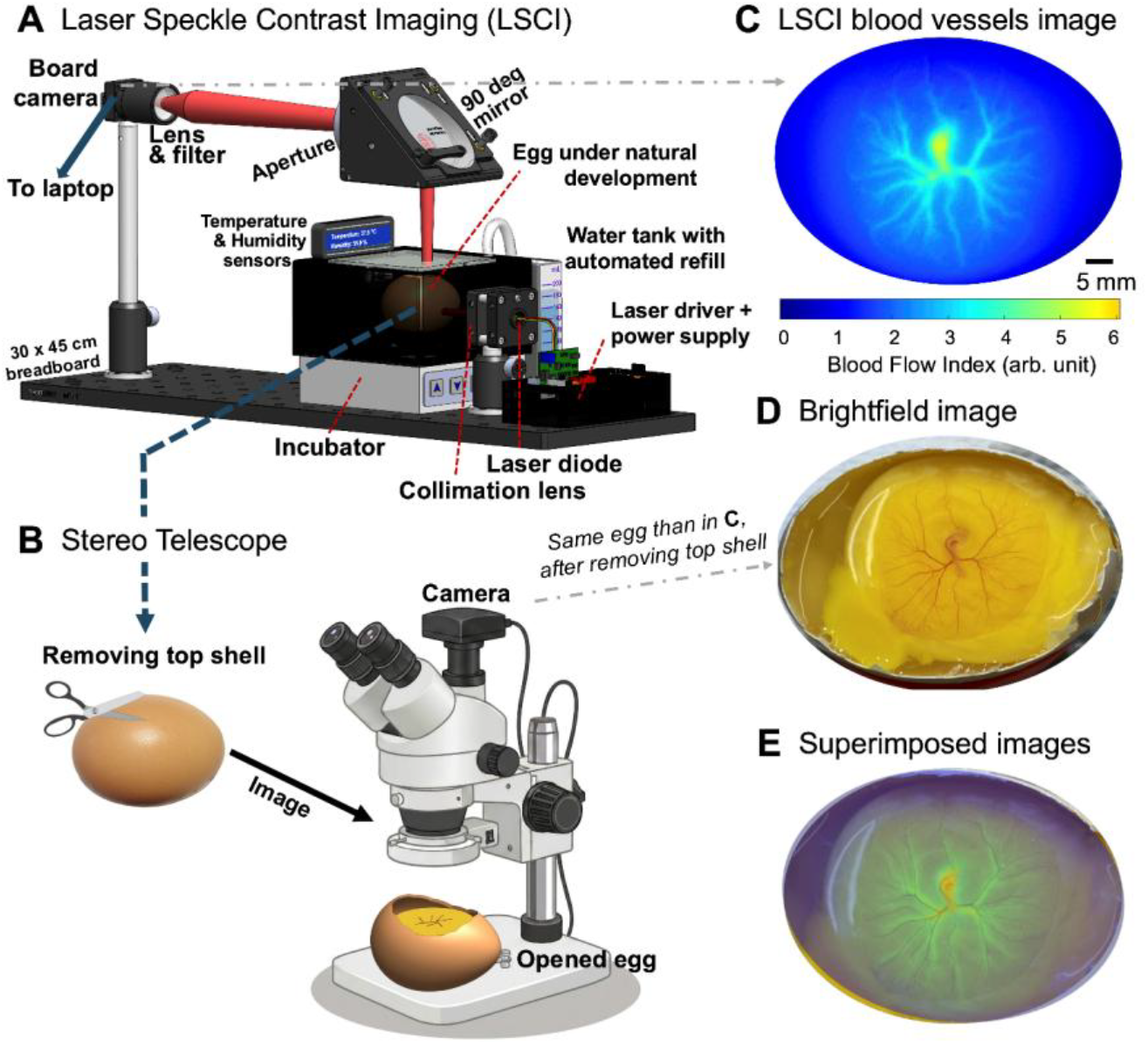
Comparison of automated laser speckle contrast imaging (LSCI) system with conventional brightfield imaging. **A**. Experimental arrangement of the automated LSCI system with a controlled incubation environment. **B**. Experimental arrangement for brightfield imaging where the top portion of the eggshell was removed to expose the embryo following LSCI imaging. **C**. Representative LSCI image of the heart and extraembryonic vasculature of a chicken egg on day 3 of incubation. **D**. Corresponding brightfield image of the same embryo shown in (D). **E**. Superimposed images of the LSCI and brightfield images, with the LSCI image overlaid on the brightfield image using transparency to show the matching blood vessel structures

As shown in Figs. 1C-1E, the vasculature observed with LSCI corresponds to those visible in the brightfield image. Although the brightfield image, acquired after shell removal, provides higher spatial resolution and more detailed visualization of the embryo body and its external vasculature, the LSCI image clearly demonstrates that the embryonic heart and its surrounding blood vessels can be accurately visualized through the intact eggshell. The embryonic heart appears as the region with the highest blood flow in the LSCI image. To further evaluate the system’s capability, we quantified the diameter of the smallest blood vessels detected by LSCI, see Smallest detectable blood vessels subsection of the Methods. Analysis from 3 eggs showed that the smallest detectable vessel diameter was 70 ± 17 μm (mean ± standard deviation), indicating that our system can measure both large vessels and finer structure such as arterioles, venules, and larger capillaries.

Together, these results demonstrate the effectiveness of our LSCI system for imaging the developing chick heart and extraembryonic vasculature. Importantly, the system integrates a fully automated incubation environment, incorporating continuous humidity and temperature monitoring, automatic water refilling (when humidity drops below threshold), and sending alerts to the operators if the humidity/temperature falls outside normal range. Such a system supports embryonic development with minimal user intervention. The laser is controlled by an imaging software and is switched-on only during image acquisitions. In addition to its imaging capabilities, our LSCI systems remain relatively affordable (≤$1,500 system cost and approximately $1,350 for laptop), compact (≤30×40×20 cm), and features an intuitive graphical user interface (GUI) that automates image acquisition while providing built-in safety and system-status notifications.

### 2.2 Longitudinal Imaging of the Chick Embryo

To evaluate the capability of the automated LSCI system to perform longitudinal and non-invasive monitoring, chick embryos were imaged every 30 minutes from day 0 to day 8 (11520 minutes) of incubation. For clarity, the results presented in Figures. 2 and 3 are limited to the period from day 0 to day 4.6 (6,720 minutes) of incubation, during which the major stages of early cardiovascular development occur. The complete longitudinal imaging datasets spanning day 0 to day 8 can be found in the Supplementary movies. A total of N=15 eggs were imaged, distributed in three groups of n=5 eggs: brown (Rhode Island), white (White Leghorn), and green (Easter Egger) egg. These breeds were selected to account for variations of eggshell color and breed diversity.

**Figure 2.**
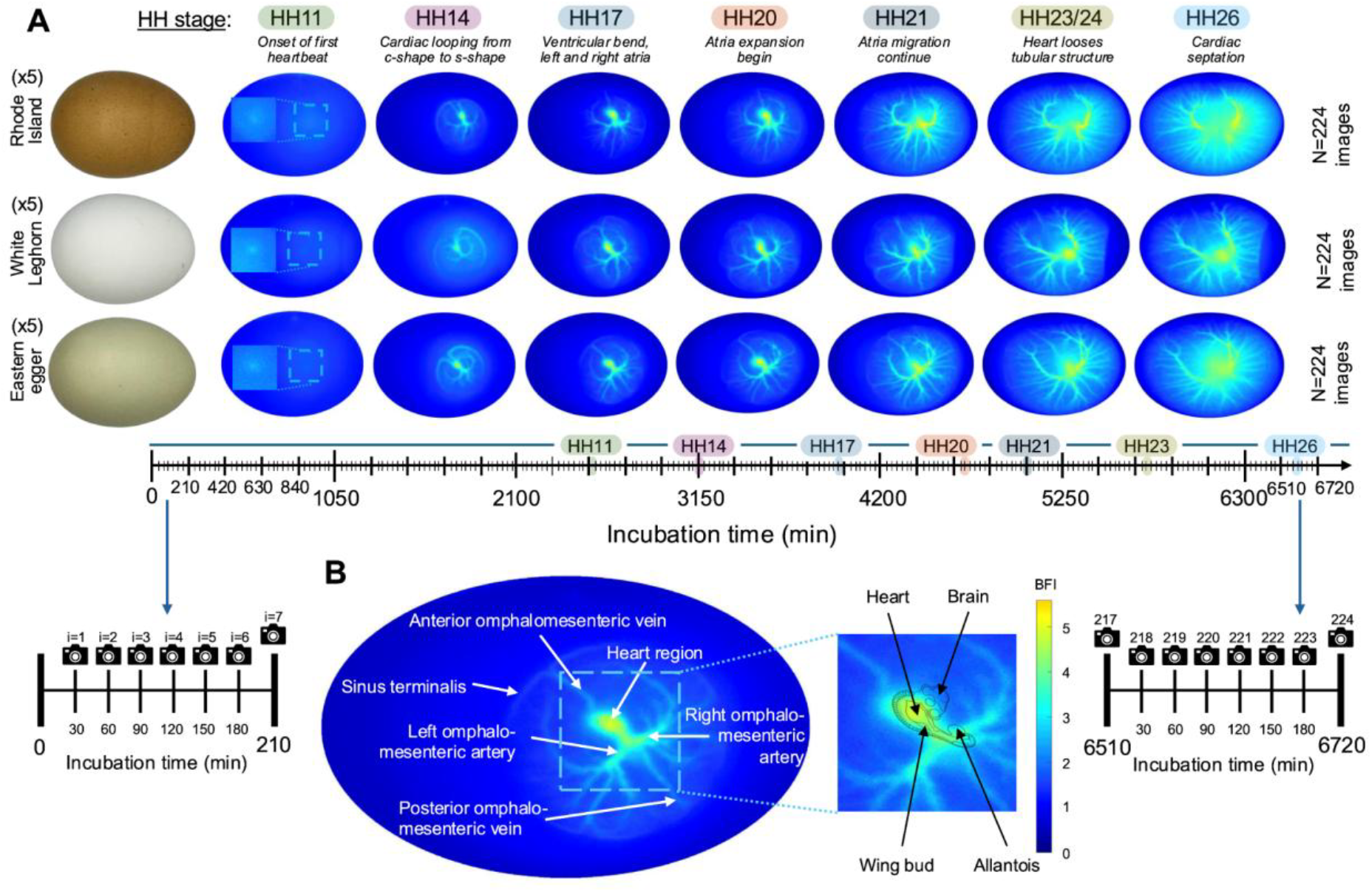
Longitudinal imaging of developing chick embryos from 0 to 6720 minutes (HH26). **A**. Selected LSCI images from three representative embryos are shown in each row. A total of 224 images were recorded for each embryo with an interval between two images of 30 minutes. **B**. LSCI image at HH20, where heart region and major arteries and veins are labeled. A time-lapse movie spanning days 0–8 can be found in the digital Supplementary Information.

**Figure 3.**
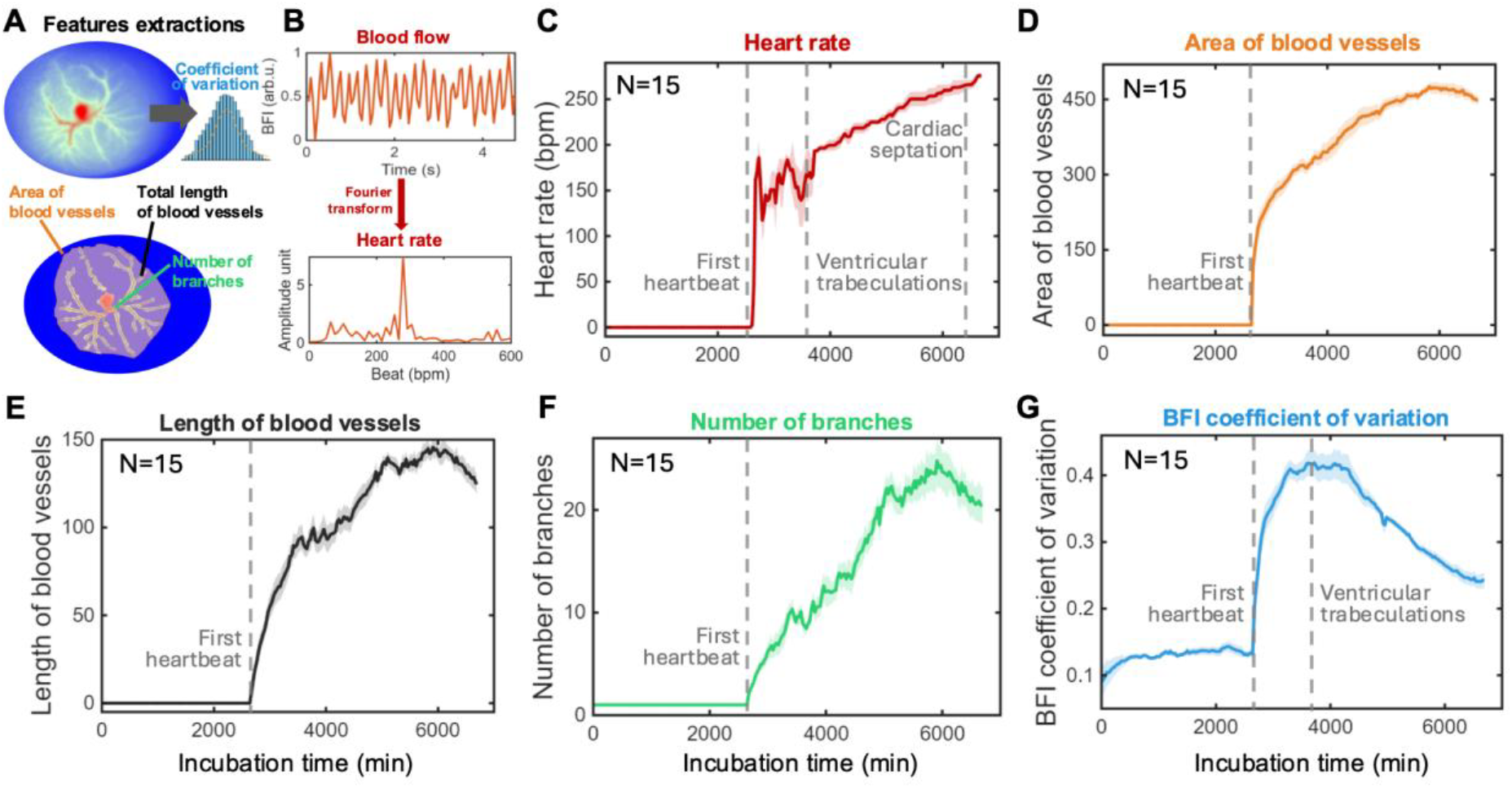
Changing cardiovascular features during embryonic development. **A**. Schematic of the extracted features from structural LSCI imaging. **B**. Schematic of the extracted heart rate from functional LSCI imaging. **C– G**. Extracted developmental features as a function of incubation time, averaged from N=15 chick embryos. **C**. Heart rate. **D**. Area of blood vessels. **E**. Total length of blood vessels. **F**. Number of branches. **G**. The blood flow index (BFI) coefficient of variation. Lines represent the mean values from N=15, and shaded regions indicate the ± standard error.

The imaging results are shown in **Figure** 2A, where the developmental time is quantified in incubation minutes. As shown, the first detectable stage is a small bright spot usually near the center of the egg, corresponding to Hamburger–Hamilton^[25]^ stage 11 (HH11), i.e. 44 hours of incubation. This corresponds to the first heartbeat of the chick embryo and the onset of cardiac angiogenesis^[26]^. Note that the timing of the first heartbeat was not identical across embryos, but for consistency and comparative analysis, we defined the first detectable heartbeat as the 2,640 minutes timepoint (i.e. stage HH11-12). This establishes a reliable baseline for characterizing vasculogenesis in the chick embryo model. One can reliably monitor embryonic development and precisely stage chick embryos by matching LSCI images to the Hamburger Hamilton reference atlas^[21]^. By approximately HH26 (6720 minutes), the vascular network has expanded to cover the entire two-dimensional field of view, after which further development is characterized primarily by three-dimensional growth, including increased vessel diameter and depth.

This is the first study to achieve continuous, non-invasive longitudinal imaging of vascular development in the same chick embryo throughout early embryogenesis.

Figure 2B demonstrates the imaging capability of our LSCI system. As shown, the heart region as well as major arteries and veins, can be visualized. Although the morphological changes of the heart from HH10 to HH26 cannot be directly resolved, the progressive growth of the heart can be inferred.

### 2.3 Quantitative analysis of cardiovascular features

Next, we analyzed five quantitative metrics to characterize chick embryo development (**Figure** 3). We first applied a feature extraction algorithm^[21,23]^ to the LSCI structural images to quantify three vascular features (Figure 3A): (1) area of blood vessels, (2) length of blood vessels and (3) number of branches. From the blood flow index (BFI) intensity distribution (LSCI image), we also calculated (4) the BFI coefficient of variation which is a measure of relative variability in BFI values, calculated as the ratio of the BFI standard deviation over the mean^[27]^. Details about the calculations are provided in the Feature extraction subsection of Methods. A unique capability of our LSCI system is its ability to non-invasively monitor blood flow and heart rate (Figure 3B) utilizing the temporal blood flow information, i.e. LSCI functional imaging. We extracted (5) heart rate for each imaging dataset. Details regarding heart rate calculations are provided in the Blood flow subsection of Methods and in^[21]^. All these five quantitative metrics are presented as a function of incubation time in Figures. 3C–G.

In Figures. 3C-G, the solid line represents the mean curve 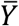 calculated from the N=15 eggs recorded (three eggs showed in Figure 2), while the shaded area represents the standard error (±*SE*), defined as:

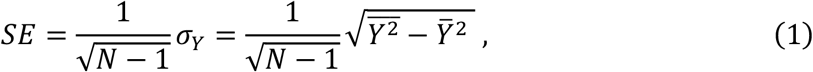

where *σ*_*Y*_ is the sample standard deviation of the feature across the N=15 embryos, and *Y* is the extracted feature (e.g. area of blood vessels).

The results for each metric are presented in Figures 3C-G. Overall, although all features display consistent developmental trajectories across embryos, the temporal patterns of growth slightly differ. Heart rate (Figure 3C) exhibits an unstable period from HH11 to HH16, which might be due to the transition of the heart from a simple linear tube to a looping structure. The heart then stabilizes and increases steadily from HH16 onward. The area (Figure 3D) and length (Figure 3E) of blood vessels exhibit similar growing curves, characterized by an initial sharp increase followed by a linear increase, indicating that both area and length continuously increase during embryonic development rather than increasing through distinct developmental stages. In contrast, the number of branches (Figure 3F) increases in a stepwise manner, with intermittent increments rather than continuous growth. Finally, the BFI coefficient of variation (Figure 3G) exhibits an initial sharp rise, followed by a plateau and subsequent decrease, a trend that can be explained by changes observed in the heart rate curve. Altogether, these features provide complementary and coherent information about embryonic development, ideal for an automated staging system. Finally, it is important to note that the standard error for all the curves remains significantly low.

From Figure 3C, one can see that from the onset of first heartbeat (HH11) to approximately 3,700 minutes of incubation (61.7 hours, HH16), the heart rate is inconsistent and fluctuates between 100 to 200 bpm (beats per minute). Then the heart rate becomes more regular and increases steadily with embryonic development, reaching about 275 bpm by 6,720 minutes of incubation (112 hours, HH26). Note that, by design, our heart rate extraction algorithm assigns a heart rate of 0 until blood vessels are first detected (see Methods), which occurs at the onset of vascularization shown in Figure 3D at 2,640 minutes (44 hours, HH11-12)

The BFI coefficient of variation quantifies the variability in blood flow across different regions of the egg, where higher value indicates that some vessels exhibit high blood flow while others exhibit low blood flow. Conversely, a lower value reflects a more uniform distribution of blood flow throughout the vascular network. In our data, before the onset of the first heartbeat (0-2,640 minutes, 0-44 hours), BFI coefficient of variation remains close to zero. Following the initiation of cardiac activity, BFI coefficient of variation increases rapidly with a significantly low standard error, indicating highly consistent measurements across embryos, until approximately 2,850 minutes of incubation time (47.5 hours, HH12). From 2,850 minutes to 3,300 minutes (47.5-55 hours), the increase becomes more gradual, reaching a plateau at around 3,300 minutes (55 hours, HH15). The BFI coefficient of variation then remains relatively stable throughout this plateau phase, from 3,300 to 4,300 min (55-71.7 hours), with a maximum observed at 3,720 minutes (62 hours, HH16), which coincides with the onset of the linear increase in heart rate (Figure 3C). Following the plateau phase, the BFI coefficient of variation gradually decreases. We hypothesize that this trend reflects the maturation of the embryonic cardiovascular system. From the first heartbeat at HH11 until the establishment of an approximately linear heartbeat trend at HH16, vascularization is not yet fully developed, i.e. some vessels exhibit high blood flow while others exhibit low blood flow, resulting in an increase in BFI coefficient of variation. Once the heartbeat is consistent and increases steadily, the vascular network is more established, and blood flow becomes more uniform across vessels. Therefore, the BFI coefficient of variation gets lower. We anticipate that extending the measurements beyond 7,000 minutes would result in a plateau state (i.e. stable BFI coefficient of variation).

### 2.4 Longitudinal imaging of the chorioallantoic membrane (CAM)

The chicken chorioallantoic membrane (CAM) is a highly vascularized extraembryonic membrane that performs vital functions during embryonic development such as gas exchange, nutrient transport, calcium mobilization, and waste removal^[28]^. The chick embryo CAM begins developing around Hamburger–Hamilton stage HH9 and progressively expands to support the increasing metabolic demands of the growing embryo. Due to its cost-effectiveness, accessibility, rapid development, and favorable ethical considerations, the CAM has become a well-established model for a wide range of preclinical applications, including drug testing, tumor growth, metastasis studies, and tissue engineering^[29–32]^.

As shown in **Figure** 4, our automated LSCI system enabled non-invasive visualization of the onset and longitudinal development of the CAM vasculature, from around 5,400 minutes of incubation (HH22) to around 10,080 minutes (HH31). Like early yolk sac blood vessel development (Figure 2), CAM vascular expansion occurred through a continuous process. However, compared with early-stage embryonic vasculature, the CAM exhibits a more complex and diverse vascular architecture, with greater variability between individual embryos and reduced inter-embryo consistency. Some of this may be due to the position of the embryo during imaging. The longitudinal observations indicate that the CAM vasculature progressively expands to occupy the whole egg surface area and becomes the dominant vascular network This represents the first demonstration of non-invasive and/or longitudinal observation of the CAM vascular development in the same embryo. It provides a powerful new tool for studying dynamic angiogenesis, vascular remodeling, and treatment responses in applications such as anti-angiogenic drug screening, cancer biology, metastasis, regenerative medicine, and tissue engineering.

**Figure 4.**
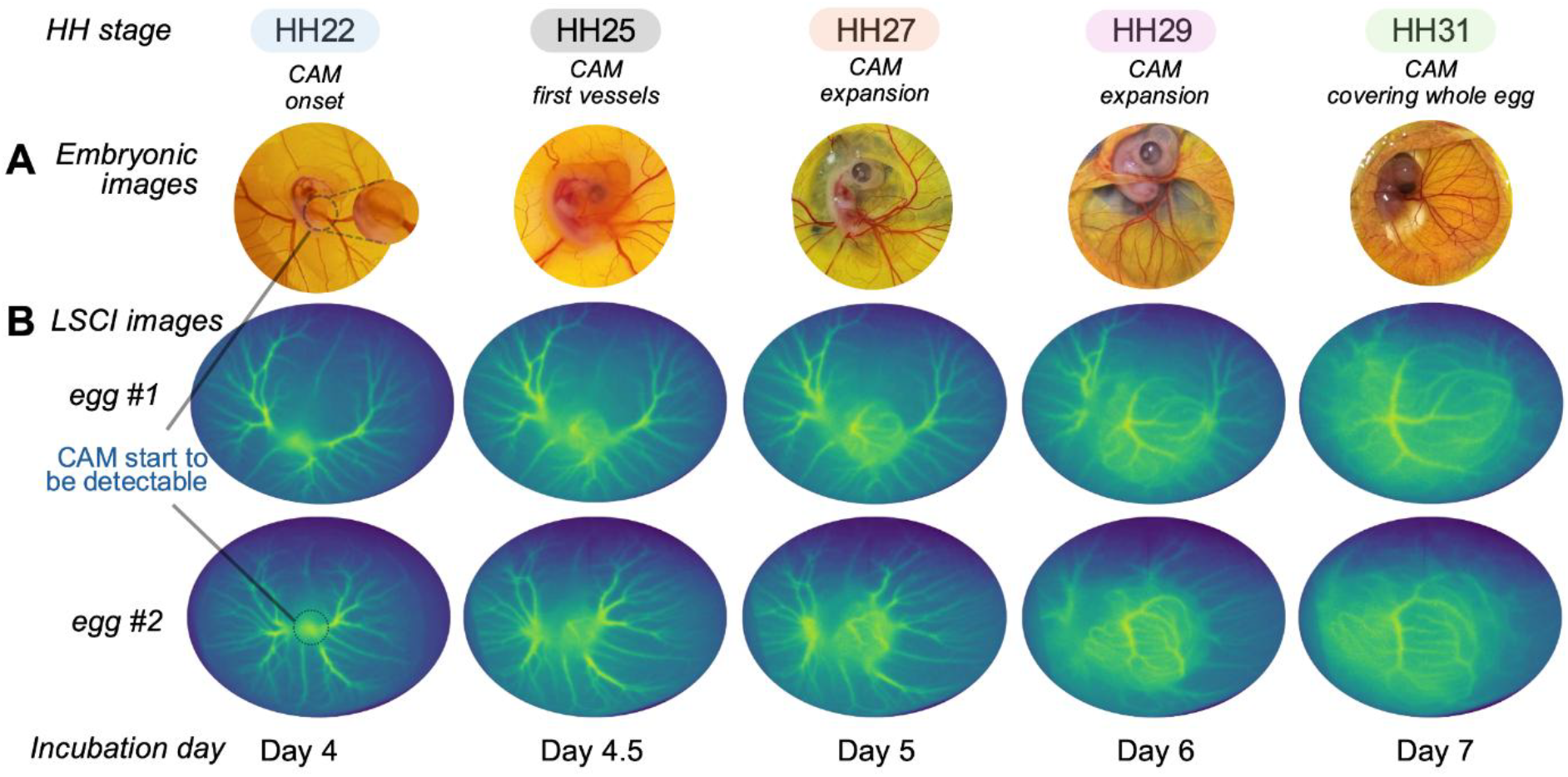
Longitudinal imaging of the chorioallantoic membrane (CAM). **A**. Typical brightfield images of the CAM after opening the eggshell. **B** and **C**. Longitudinal LSCI images depicting the onset and growing of the CAM vasculature for two different eggs. A time-lapse movie spanning days 0–8 can be found in the digital Supplementary Information.

## 3. Discussion

In this study, we developed and validated an automated laser speckle contrast imaging (LSCI) platform for continuous, non-invasive, longitudinal monitoring of extraembryonic vascular development in the living chick embryo. Most imaging approaches require egg windowing, embryo removal, labeling, fixation, or destructive sampling at discrete time points, whereas our system enables repeated measurements of the same embryo throughout development while maintaining a controlled incubation environment. The platform provides a robust and user-friendly approach (GUI interface) for long-term developmental imaging, both at a relatively affordable cost (< $1,500) and with a compact footprint (≤30×40×20 cm).

Using this system, we achieved non-invasive longitudinal visualization of chick embryonic cardiovascular development and CAM vascular expansion within the same embryo. The high temporal resolution (e.g., 30-minute imaging intervals, adjustable to specific experimental requirements) revealed dynamic vascular growth patterns that are difficult to capture using traditional endpoint-based methods. The common trajectory of early cardiac and vascular morphogenesis across breeds and shell types highlights the potential of the system for automated developmental staging. Beyond early cardiovascular development, longitudinal CAM imaging demonstrated the ability to monitor large-scale vascular remodeling during later embryogenesis. The CAM has been used extensively to study metastases, cancer biology, as well as vascular development and vasculogenesis^[33,34]^. The LSCI imaging platform provides an opportunity to study these processes *in vivo*.

The ability to repeatedly visualize vascular development in the same embryo opens new opportunities for quantitative developmental biology. One promising research direction is the development of an automated embryo staging system that integrates structural and functional imaging of embryonic hemodynamics with deep learning-based classification algorithms for high accuracy staging tasks. We hypothesize that continuous physiological measurements— including heart rate, blood flow dynamics, vascular branching patterns, vessel growth trajectories, and cardiac morphology—contain sufficient information to accurately determine developmental stage and predict developmental progression from a single longitudinal LSCI dataset, all non-invasively. The development and validation of such an automated staging platform add to the current paradigm of embryo classification, which relies primarily on invasive morphological assessment using the Hamburger–Hamilton staging system into a fully non-invasive, objective, and reproducible approach that preserves natural embryonic development. A second direction can be embryo viability assessment for the conservation and management of rare and endangered avian species, where the automated LSCI system could alert of slow or unnormal embryonic development. Another promising direction is the creation of a comprehensive developmental atlas of functional cardiac physiology and vascular morphogenesis across breeds and shell types.

Finally, the LSCI live imaging provides a platform for the study of the dynamics of heart formation and vasculogensis and for testing internal or environmental agents that might perturbate this process.

## 4. Methods

### 4.1 Laser Speckle Contrast Imaging (LSCI)

Laser speckle contrast imaging (LSCI) utilizes laser speckles originating from the scattering of coherent laser light by biological tissue (sample)^[35–38]^. Laser speckles arise from the constructive and destructive interference of the scattered light with itself^[38]^. As dynamic structures within biological tissue move, the speckle pattern undergoes temporal fluctuations. For example, the motion of red blood cells within blood vessels causes temporal variations in the speckle intensity, which can be quantified to estimate blood flow. By recording the speckle pattern with a camera and analyzing its temporal fluctuations across successive image frames, the spatial distribution of blood flow within the sample can be reconstructed. For that, speckle contrast is calculated from the camera images as:

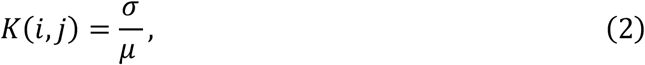

Where K is the speckle contrast at pixel (i,j), *σ* is the standard deviation, and *μ* is the mean. As the velocity of red blood cells increases, the speckle pattern decorrelates more rapidly, resulting in a lower speckle contrast. Conversely, slower blood flow produces a higher speckle contrast. A more detailed description of the LSCI principle for egg imaging can be found in ^[21]^.

In our system, we employ temporal LSCI^[39,40]^, where the camera recorded a sequence of N = 100 speckle frames images when the laser was on and another 100 noise frames when the laser was blocked with the same exposure time T to generate noise subtracted speckle frames Ĩ(t) to reduce noise. The noise frames are then averaged to a single frame and subtracted from the speckle frames^[21]^. The temporal speckle contrast *K*_*t*_ was calculated for each camera pixel Ĩ(*i, j*; *t*) in the temporal domain over the N noise subtracted speckle frames as:

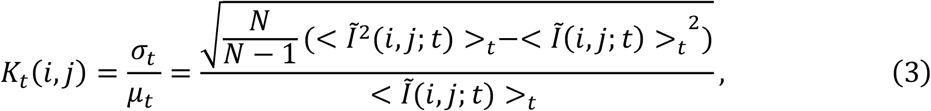

where *Ĩ*(*i, j*) is the intensity at the pixel row *i* and column *j, t* is the time at which the speckle pattern was recorded by the camera, and <>_*t*_ indicates temporal averaging over time occurring at pixel *i, j*. The speckle contrast corresponds to the ratio between the standard deviation *σ*_*t*_ and the temporal mean *μ*_*t*_ of the N recorded speckle pattern images. The blood flow index (BFI) was calculated from the speckle contrast as ^[41]^:

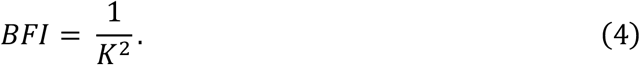

Blood flow index (BFI) provides a relative measure of blood perfusion, reflecting changes in the amount of blood flowing through the tissue over time. Variations in the BFI may result from changes in blood flow velocity, vessel diameter, or both. Note that blood flow index (BFI) and blood volume index (BVI) are related; however, their relationship between the two from speckle imaging remains a subject of ongoing investigation^[42,43]^. **Figure** 1B presents a representative image of an egg’s vascular network reconstructed using temporal LSCI. The vascular network is clearly visualized, encompassing vessels of varying diameters.

### 4.2 Smallest detectable blood vessels

To evaluate the imaging performance of our LSCI system, we quantified the diameter of the smallest blood vessel that could be detected from three eggs. For each egg, an LSCI image was first acquired. Then, the top eggshell was removed, and the corresponding vascular region was imaged using bright-field microscopy to directly visualize the blood vessels, **Figure 5**A. During bright-field imaging, a calibration target was placed on top of the embryo to determine the pixel-to-distance conversion factor (μm/pixel), Figure 5A. This calibration established scalebar, allowing smallest blood vessel detected in LSCI images to be converted into their corresponding physical dimensions (diameter, μm). For that, a line profile perpendicular to the vessel axis was extracted and the vessel diameter was determined using the full width at half maximum (FWHM)of the intensity profile. The FWHM value, measured in pixels, was converted to the corresponding physical vessel diameter (μm) using the pixel-to-distance conversion factor. This procedure was repeated over three blood vessels for three eggs, and a mean ± standard error was calculated to characterize the vessel detection capability of our LSCI system, as shown in Figure 5B.

**Figure 5.**
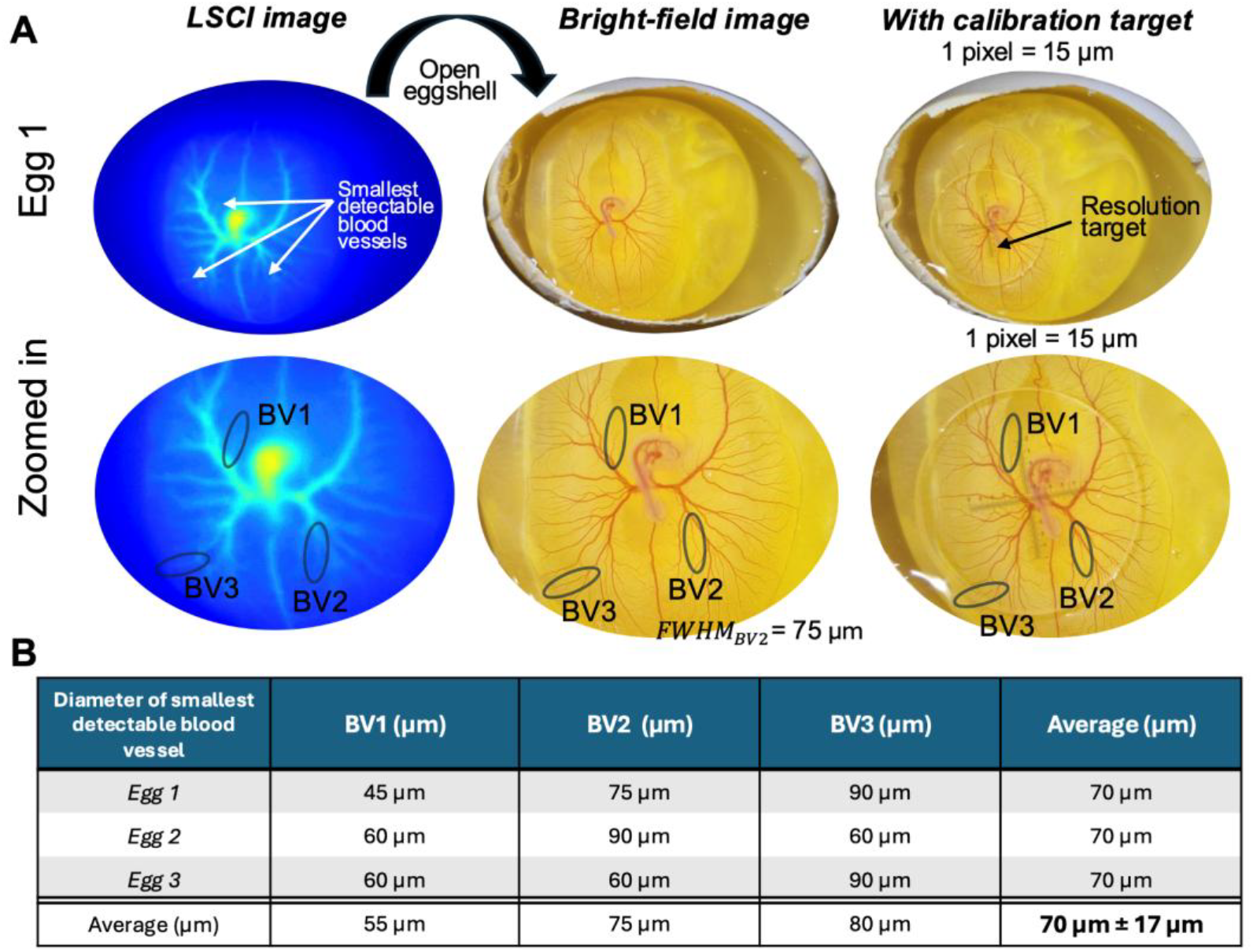
Smallest detectable blood vessels in LSCI system. **A**. Three images of the same egg acquired at the same time: an LSCI image, a bright-field image, and a bright-field image with a calibration target. Three smallest detectable blood vessels in LSCI were labeled. **B**. Diameter of the three smallest detectable blood vessels for three individual eggs with a total average value of 70 ± 17 μm determining the vessel detection capability of our LSCI system.

### 4.3 Extracting blood flow dynamics

We previously demonstrated that blood flow dynamics in chicken eggs can be extracted from LSCI recordings^[21,23]^. Rather than computing temporal LSCI from N = 100 consecutive speckle frames, blood flow dynamics can be monitored by applying a sliding window of n = 3 adjacent frames to generate a time-varying temporal speckle contrast. The resulting contrast values are subsequently converted into relative blood flow values (BFI) over time. The blood flow extraction algorithm is described in detail in^[21]^ and has been shown to produce results comparable to those obtained using the speckle visibility spectroscopy (SVS, also named speckle contrast optical spectroscopy SCOS) method^[41,44]^. Specifically, both methods yielded nearly identical heart rate measurements and blood flow dynamics when applied to the same chicken egg LSCI dataset. In the SVS/SCOS approach, blood flow is estimated from the spatial speckle contrast of a single speckle image, with each camera pixel serving as an independent sampling point.

### 4.4 Heart rate calculations

As explained in the previous section, it is possible to measure the blood flow of a chicken egg using our LSCI system enables measurement of blood flow in chick embryos (Figure 3B). During calculation, the speckle frame was cropped into a rectangular region of interest (ROI). The ROI encompassed both the heart region (brightest peak) and the embryo blood vessels. The same ROI was applied to all eggs, regardless of developmental stage. Ideally the ROI should be cropped specifically for each developmental stage to the size of the embryo and the surrounding blood vessels.

To extract heart rate, Fourier transform of the BFI was computed (Figure 3B, bottom part), and the prominent peak in the frequency spectrum was identified as the heart rate frequency (Hz). The frequency was subsequently converted to beats per minute (BPM). In our measurements, we use N=100 frames and recording speed of *fps* = 21 frames per second. In Fourier analysis, the frequency resolution is defined as Δ*f* = 1/*T* where *T* is the total duration of recording. In our case, *T* = *N*/*fps* = 4.8 seconds, leading to Δ*f* = 0.21 Hz = 12.5 bpm. This sets our standard error on the heart rate (HR) measurement to be HR ± Δ*f*/2 = 6.25 bpm. We acknowledge this standard error could be reduced by increasing the recording duration.

For each LSCI recording, heart rate was extracted as follows: As embryonic heart rate changes gradually within the 30-min interval, the heart rate detected at the previous time point was used as a reference for subsequent measurements. Specifically, the dominant frequency peak is restricted to a narrow window extending from -1.5 Hz to +2.5 Hz relative to the previously detected heart rate. We assume the heart rate to be 0 when *t* = 0. This approach reduced the likelihood of incorrectly identifying noise-induced peaks as cardiac signals, especially when the heart is beating inconsistently. Before the formation of vascular network, i.e. where no blood vessels were detected (area of blood vessels = 0), the corresponding heart rate measurement was assigned a value of 0.

### 4.5 Automated LSCI setup

Figure 1A depicts the experimental setup of our automated LSCI system. A coherent laser diode (Thorlabs L852SEV1) operating at a wavelength of 852 nm served as the illumination source. The laser output power was set to approximately 250 mW and remotely controlled. The laser was driven by a Thorlabs EK1101 laser driver board interfaced by a custom-designed printed circuit board (PCB). The PCB was powered by a 9 V battery, which provides several hours of operation. The laser beam was directed into the incubator chamber to illuminate the egg. Note that the laser was collimated using an aspherical lens of focal length f=11 mm, numerical aperture 0.26, and anti-reflective coating 650 - 1050 nm (Thorlabs A220TM-B).

The egg was housed within a custom-designed, 3D-printed incubator chamber. The chamber was mounted on an incubator heating plate which incorporates a small fan to ensure uniform circulation of warm air. The heater was regulated to maintain an internal chamber temperature of 37.5°C, providing optimal conditions for embryonic development. The chamber includes a laser entrance window sealed with transparent tape to allow the incident laser beam to illuminate the egg. A 3-mm-thick plexiglass window was positioned above the egg for the scattered light to be collected by the camera. Environmental conditions inside the chamber were continuously monitored using a temperature and humidity sensor (Adafruit SHT31) interfaced with a microcontroller (Arduino R4 WiFi). If the measured temperature fell below 35°C, the system sent a warning notification to the operator. If the humidity level dropped below 50%, the Arduino triggered a submersible pump, dispensing approximately 15 mL of distilled water into the chamber. If the humidity remained below 50% after refilling, the system sent a warning notification to the operator. The automated system maintained stable incubation conditions with minimal user intervention, reducing the risk. The custom-built incubator consistently achieved higher embryo development rates compared with commercially available 12-egg and 48-egg incubators. Compared to previous incubation systems^[21,23,24]^, the key innovation of our design is that the egg is housed directly within the incubator chamber and has automated refill functions.

The scattered light exiting the egg from the top surface was redirected onto a camera using a 90-degree mirror (Thorlabs PFE20-P01 and KCB2EC). The camera was a monochromatic camera (Thorlabs CS126MU) of 3000 × 4096 pixels, with a pixel pitch size of 3.45 × 3.45 μm. The exposure time was set to 10 ms and at a speed of 21 frame per second (FPS). In front of the camera, a lens of 50 mm focal length (Edmund Optics #86-574) was positioned to focus the camera onto the chick embryo plane. An aperture located near mirror mount plane, the imaging lens aperture, and the camera distance from the egg were jointly optimized to provide a field of view encompassing the entire egg while ensuring that the speckle size at the camera sensor exceeded two pixels, satisfying the sampling criterion for accurate speckle contrast measurements^[45–47]^. Thus, in our optical setup, the system magnification was 0.2, and the numerical aperture was 0.03. The LSCI recording code and algorithm were as described in^[21]^. Note that the laser was only switched-on during recording period. The LSCI system was enclosed in a black box to prevent laser light leakage outside the optical setup and to prevent room and stray light noises.

### 4.6 Animal study

The animal research for this study received confirmation from the Institution Animal Care and Use Committee (IACUC) at Stevens Institute of Technology. In all experiments, chicken eggs were not incubated later than day 12. As such, the IACUC exempted this animal study from an IACUC protocol. All the experiments including the disposal of the chicken egg embryo were performed in compliance with Stevens IACUC policies. Fertilized chicken eggs were obtained from Bone-In Food, New Jersey, a farm cooperative. To account for biological variability, eggs were typically sourced from three different locations. The eggs were incubated in commercial incubator base Hethya, HHD and with our custom-built incubator chamber. The incubators automatically maintained the temperature between 37.5–39.0°C, while the relative humidity was regulated within the 50–85% range. During incubation, the eggs were not turned for longitudinal imaging experiments (standard incubation protocols recommend turning them every 60–90 min). This did not affect the development rate of embryos.

### 4.7 Feature extraction

To apply feature extractions, the LSCI blood-flow images were processed using a custom MATLAB-based image-processing method, detailed in^[21]^. First, the eggshell region was detected. An inner elliptical mask was then created, with its major and minor axes set to 75% of the detected eggshell dimensions. This helped reduce noise and unwanted structures near the egg boundary. Then, the image inside the elliptical region was enhanced using filtering, contrast improvement, background subtraction, and smoothing, as detailed in^[21]^. Automatic thresholding was then used to separate the blood vessels from the background and create a binary vessel mask. Small regions were removed, and gaps between nearby vessel regions were reduced. The cleaned vessel mask was then converted into a one-pixel-wide skeleton. Short false branches and peripheral ring-like structures were removed, and branch points were detected.

Four features were extracted from each image: area of blood vessels, length of blood vessels, number of branches, and BFI coefficient of variation. Area of blood vessels was calculated from the number of pixels in the cleaned vessel mask. It represents the total image area occupied by the segmented blood vessels. A larger vessel area indicates greater spatial coverage of the vascular network. Length of blood vessels was calculated from the total length (one-dimensional skeleton distance) of the detected blood vessels. A higher vessel-length value indicates that the vascular network extends over a greater distance. The number of branches was calculated using the detected branch points and connected vessel regions. It represents the structural complexity of the vascular network. A higher number of branches indicates that the vessels are more divided and interconnected. For each image, the BFI coefficient of variation was calculated and normalized from the distribution of BFI values within the detected vessel region as the ratio of the standard deviation to the mean. A low value indicates that the blood-flow-related values are relatively uniform, whereas a higher value indicates greater variation across the vascular network.

## Acknowledgements

This work was supported by startup funds provided to S.M. by Stevens Institute of Technology.

## Data Availability Statement

The datasets generated during and/or analyzed during the current study are available from the corresponding author on request.

## Supporting Information

### Movie S1

Longitudinal imaging of extraembryonic vascular development in the intact eggshell from day 0 to day 8 of incubation. The video visualizes the dynamic progression of embryonic development while preserving its natural developmental environment, with text highlighting key developmental stages and vascular changes throughout incubation.

### Movie S2

Unannotated original video of the longitudinal imaging described in Movie S1, showing extraembryonic vascular development from day 0 to day 8 of incubation.

Movies S1 and S2 can be found from this link: https://drive.google.com/drive/folders/1Am7npE9HqNHRRoXGeNldNjAVzrCEouOz?usp=drive_link

